# A shared linguistic space for transmitting our thoughts from brain to brain in natural conversations

**DOI:** 10.1101/2023.06.27.546708

**Authors:** Zaid Zada, Ariel Goldstein, Sebastian Michelmann, Erez Simony, Amy Price, Liat Hasenfratz, Emily Barham, Asieh Zadbood, Werner Doyle, Daniel Friedman, Patricia Dugan, Lucia Melloni, Sasha Devore, Adeen Flinker, Orrin Devinsky, Samuel A. Nastase, Uri Hasson

**Author notes:** These authors contributed equally to this work (Samuel A. Nastase, Uri Hasson).

## Abstract

Effective communication hinges on a mutual understanding of word meaning in different contexts. The embedding space learned by large language models can serve as an explicit model of the shared, context-rich meaning space humans use to communicate their thoughts. We recorded brain activity using electrocorticography during spontaneous, face-to-face conversations in five pairs of epilepsy patients. We demonstrate that the linguistic embedding space can capture the linguistic content of word-by-word neural alignment between speaker and listener. Linguistic content emerged in the speaker’s brain before word articulation, and the same linguistic content rapidly reemerged in the listener’s brain after word articulation. These findings establish a computational framework to study how human brains transmit their thoughts to one another in real-world contexts.

## Introduction

Language is the bedrock of human communication, allowing us to share our ideas and feelings with others. Successful communication, however, relies on a shared agreement regarding the meaning of words *in context*. For example, the word *cold* can describe the temperature, a personality trait, or a viral infection, depending on the context. This contextual meaning of language resides in a shared space between people in a community of speakers: words absorb transient, agreed-upon meanings specific to their use and context (Wittgenstein, 1953; Dor, 2015). Without a shared agreement, it would be impossible for strangers to understand one another. For example, speakers can only understand whether the word *cold* in the sentence “you’re as cold as ice” refers to a personality trait or physical temperature if they are aware of the conversational context.

Until recently, we lacked a precise computational framework for modeling how humans use words in context as we communicate with others. To overcome this limitation, previous studies of the neural basis of communication have resorted to measuring direct coupling or alignment between brains by using the neural activity in the speaker’s brain to make model-free predictions of the listener’s brain activity. These studies have revealed that stronger speaker–listener neural coupling correlates with more successful communication (Stephens et al., 2010; Hasson et al., 2012; Nastase et al., 2019; Meshulam et al., 2021; Nguyen et al., 2022; Davidesco et al., 2023). Although these analyses can quantify the strength of brain-to-brain coupling, they are content-agnostic and cannot model how we use words in context to convey our thoughts to others.

A new class of large language models (LLMs) has recently emerged that, for the first time, respects the richness of context in natural communication. Remarkably, these models learn from much the same shared space as humans: from real-world language generated by humans. LLMs rely on a simple self-supervised objective (e.g. next-word prediction) to learn to produce context-specific linguistic outputs from real-world corpora—and, in the process, implicitly encode the statistical structure of natural language into a multidimensional embedding space (Manning et al., 2020; Linzen & Baroni, 2021; Pavlick, 2022). The capacity of these models to generate fluent, context-rich text, engage in dialogue, and meaningfully answer questions is a testament to just how much can be learned from the shared space of language and communication (Piantadosi, 2023). Interestingly, recent studies have suggested that LLMs and the brain converge on shared computational principles for natural language comprehension (Heilbron et al., 2020; Schrimpf et al., 2021; Goldstein, Zada, et al., 2022; Caucheteux & King, 2022; Kumar et al., 2022).

In this study, we positioned LLM contextual embeddings as an explicit model of the shared linguistic space by which a speaker communicates their thoughts—i.e., transmits their brain activity—to a listener in natural contexts. We recorded cortical activity using electrocorticography (ECoG) in five dyadic pairs of epilepsy patients during spontaneous, interactive conversations. Modeling speech production and comprehension in this free-form setting is challenging: speakers are not constrained to a particular vocabulary or fixed turns; they are free to interrupt each other and articulate whatever words they want whenever they want. Each conversation was unique. We extracted contextual embeddings for each word in the conversation from a widely-used contextual language model, GPT-2 (Radford et al., 2019). We then trained encoding models to predict brain activity during speech production and comprehension in held-out segments of the conversations. We first demonstrate that contextual embeddings, with high temporal specificity, can predict neural activity across the cortical language network during speech comprehension and speech production. Consistent with the flow of information during communication, we use an intersubject speaker–listener encoding analysis to demonstrate that the same linguistic content in the speaker’s brain before word articulation re-emerges, word-by-word, in the listener’s brain after each word is spoken. This demonstrates that speaker and listener neural responses during natural conversations are coupled to a shared linguistic space and suggests that LLMs provide a novel computational framework for studying how we transmit our thoughts to others.

## Results

We recorded cortical activity using ECoG in five dyadic pairs of epilepsy patients during free-form, face-to-face conversations. We filtered the neural data to high-gamma broadband activity to approximate local field potentials. For each dyad, we spliced the neural data into word-level epochs, collated these epochs according to speaker and listener roles, and split the collated data for 10-fold cross-validation. We used time-resolved transcriptions of each conversation to extract embeddings for each word from the autoregressive large language model GPT-2 (Radford et al., 2019). We used ridge regression to estimate separate encoding models for the speaker and listener to predict the high-gamma band neural activity for each word using the embeddings from GPT-2 (Fig. 1A) (Holdgraf et al., 2017). To focus on electrodes that encode linguistic content, we performed a permutation test by re-estimating within-subject production and comprehension encoding models on phase-randomized neural data (*p* < .01, FDR corrected); these models were estimated from non-contextual, static GPT-2 embeddings to minimize selection bias for the contextual embeddings.

**Fig. 1.**
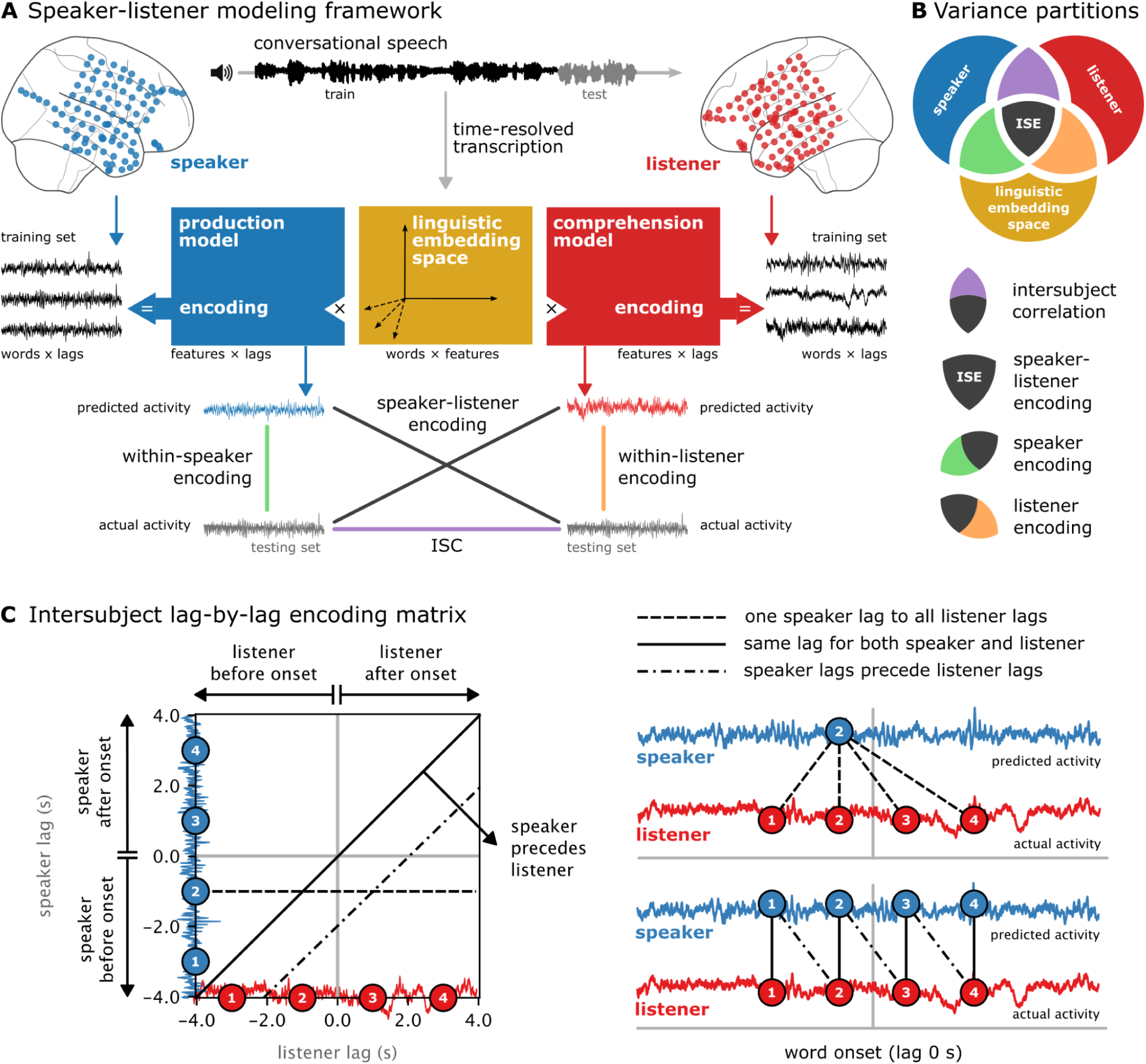
Encoding models for capturing speaker–listener linguistic coupling. (**A**) Schematic depicting how encoding models isolate the linguistic content of speaker–listener brain-to-brain coupling. Word-level neural signals for both patients are collated into speaker–listener roles and split into temporally contiguous train/test folds for cross-validation, along with their respective word embeddings. Electrode-wise encoding models are estimated using ridge regression separately for the speaker (blue weight matrix) and listener (red weight matrix) to predict neural activity from the LLM embeddings at varying lags relative to word onset. Within-subject encoding performance is quantified separately for the speaker and listener as the correlation between predicted and actual neural activity for left-out test segments of each conversation. Intersubject speaker–listener encoding is quantified as the correlation between the model-predicted activity from the speaker (or listener) and the actual neural activity of the listener (or speaker). While intersubject correlation (ISC) directly correlates brain activity between speaker and listener (purple line), intersubject encoding (ISE) effectively isolates the linguistic component of brain-to-brain coupling (black lines). (**B**) Components of speaker–listener brain-to-brain coupling. Each circle represents the variance due to one of three sources: the speaker’s brain activity (blue), the listener’s brain activity (red), and the linguistic embedding space (yellow). Labeled intersections below represent partitions of shared variance: direct intersubject correlation capturing shared variance between the speaker and listener (purple) as well as shared variance between the contextual embedding space and the speaker’s (green) and listener’s (orange) brain activity. The central intersection of all three sources (black) represents the shared variance in speaker–listener brain-to-brain coupling captured by the contextual embeddings, i.e., intersubject speaker–listener encoding (ISE). (**C**) Intersubject encoding performance is evaluated at each pair of lags between the speaker’s and listener’s brain activity, resulting in a lag-by-lag encoding matrix of correlation values (left). Word onset is depicted with gray lines at the center, dividing the matrix into quadrants: e.g., the bottom right quadrant corresponds to the speaker’s brain activity before word onset and the listener’s brain activity after word onset. The horizontal dashed line connects the speaker’s lag labeled “2” before word onset to all lags in the listener—depicted graphically on the right using dashed lines. The solid diagonal of the matrix corresponds to simultaneous speaker and listener lags, schematically shown at the bottom right as vertical solid lines between speaker–listener lags 1–4. Lag pairs under the diagonal indicate that the speaker precedes the listener (dash-dot lines), connecting speaker lags 1–3 to listener lags 2–4.

### Contextual embeddings predict brain activity in both the speaker and listener

We assessed whether the linguistic embedding space could capture time-resolved, word-related neural activity in both the speaker and the listener. We trained encoding models with 10-fold consecutive cross-validation at lags ranging from –4 s to +4 s relative to word onset and measured the correlation between actual and predicted word-related activity in each test fold separately for speech production and speech comprehension (Fig. 2). During speech production, we found maximal encoding performance in speech articulation areas along the precentral motor cortex, in the superior temporal cortex, and in higher-order language areas in the temporal pole, inferior frontal gyrus, and supramarginal gyrus (Fig. 2A, blue). Detailed inspection of the encoding performance within the speaker’s brain at various lags revealed that the maximal prediction peaked ∼300 ms before word onset (Fig. 2B, blue). During speech comprehension, the encoding model predicted neural responses in similar brain areas, particularly the superior temporal cortex (Fig. 2A, red). Comprehension encoding performance increased gradually at word onset in superior and anterior temporal electrodes, peaked ∼250 ms after word onset, and decreased over 1-second post-word-onset (Fig. 2B, red).

**Fig. 2.**
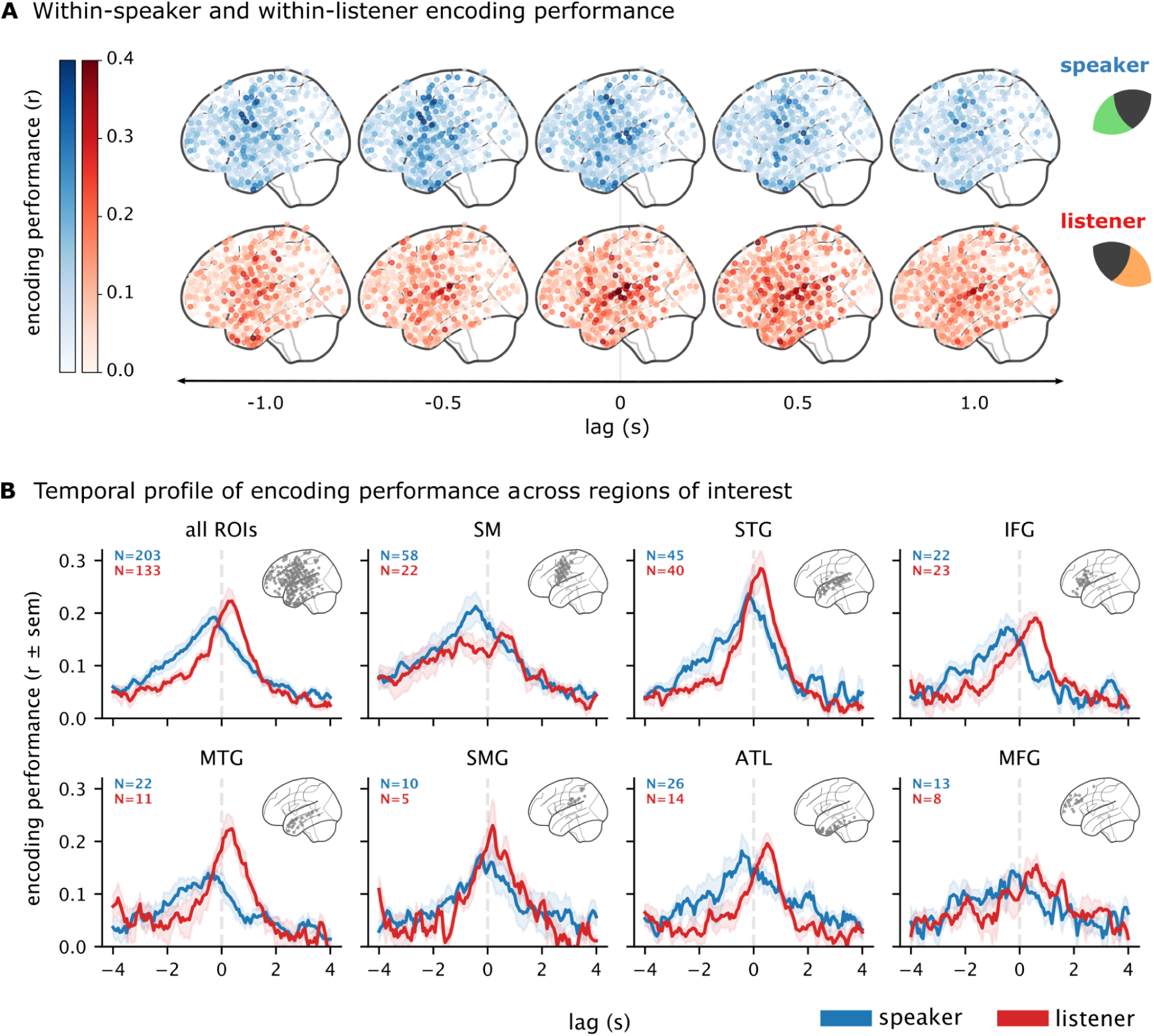
Within-speaker and within-listener linguistic encoding performance. Encoding models were trained to predict the neural activity from linguistic embeddings separately per lag and per electrode and evaluated using 10-fold cross-validation. Encoding performance is quantified as the correlation between word-by-word model-predicted and actual electrode activity. (**A**) Encoding performance for all electrodes from all subjects at five different lags relative to word onset (lag 0). Separate models are trained for spoken words (production, blue) and heard words (comprehension, red). (**B**) Encoding performance for all electrodes selected for significance (*p* < .01, permutation test, FDR corrected; see “Electrode selection” in Materials and Methods) across lags for all regions (panel B, top left) and in different regions of interest (ROIs) in the cortical language network. Error bands indicate the standard error of the mean correlation across electrodes and subjects. The number of significant electrodes in the speaker and listener is displayed in the upper left corner of each panel. SM: somatomotor cortex; STG: superior temporal gyrus; IFG: inferior frontal gyrus; MTG: middle temporal gyrus; SMG: supramarginal gyrus; ATL: anterior temporal lobe; MFG: middle frontal gyrus.

The linguistic embedding space predicts neural activity in multiple regions with different temporal dynamics and selectivity (Fig. 2B). Somatomotor (SM) electrodes encode linguistic content more so during speech production than comprehension, particularly before word articulation. Electrodes in the superior temporal gyrus (STG), on the other hand, encode linguistic content during both processes, although encoding performance is stronger for comprehension. In the inferior frontal gyrus (IFG) and anterior temporal lobe (ATL), linguistic encoding during speech production peaks prior to word onset, with sustained encoding after word articulation during comprehension. During speech production, encoding performance in most regions decreases rapidly after word onset; this decrease accompanies the increase in post-word-onset encoding performance in the listener (Table S1). These results demonstrate that the linguistic embedding space learned by GPT-2 captures relevant features for predicting neural activity during language production and comprehension across the cortical language network.

### Contextual embeddings capture linguistic coupling between the speaker and the listener

How are the speaker and listener’s brains aligned during the conversation? The previous analysis used encoding models to predict the neural signal from word embeddings separately in the speaker and listener. To assess linguistic coupling across brains, we used the encoding model trained on the speaker’s brain activity to predict the listener’s brain activity (and vice versa) using the same cross-validation scheme; that is, we correlated the model-based predictions from one brain with the actual neural activity in the other brain (Fig. 1A). This novel “intersubject encoding” (ISE) analysis quantifies how well the model fit for speech production (or comprehension) generalizes to speech comprehension (or production) in left-out segments of each conversation. By virtue of using word embeddings from a language model, the encoding model filters out non-linguistic features that may be common between the conversants but that are not present in the conversation transcript. In order to capture the temporal dynamics of speaker–listener coupling, we applied this procedure for each pair of lags in the speaker and listener’s brain activity, resulting in a lag-by-lag encoding matrix of correlation values where the y-axis indexes lag in the speaker’s brain and the x-axis indexes lags in the listener’s brain, relative to word onset (Fig. 1C). In this matrix, the central axis lines signify the onset of word articulation (lag 0 s), while the matrix diagonal corresponds to simultaneous lags between speaker and listener. Intersubject encoding below the diagonal indicates that linguistic content encoded in the speaker’s brain precedes the same linguistic content encoded in the listener’s brain (Fig. 1E).

We first applied this procedure across all electrodes selected for significance to estimate overall linguistic coupling across the cortical language network (*n* = 203 production; 133 comprehensions): we averaged the model-predicted neural activity across electrodes (e.g., in the speaker) and correlated this with the averaged actual neural activity across electrodes (e.g., in the listener). We observed time-locked, speaker–listener linguistic coupling centered around the moment of articulation of each word in the conversation (Fig. 3A; training and testing on speaker or listener yielded very similar results; Fig. S1). Consistent with the flow of information during communication, the linguistic coupling falls under the diagonal, indicating that the speaker’s brain is “leading” the listener’s brain (see Fig. S2 for significance test). We found that the linguistic features of the speaker’s brain activity ∼425 ms before word onset best predicted brain activity in the listener ∼125 ms after the word was articulated. That is, linguistic content emerges in the speaker’s brain prior to the spontaneous articulation of each word, then re-emerges in the listener’s brain after each word is heard. This temporal dynamic proceeds word by word and is specific to the current word and conversation.

**Fig. 3.**
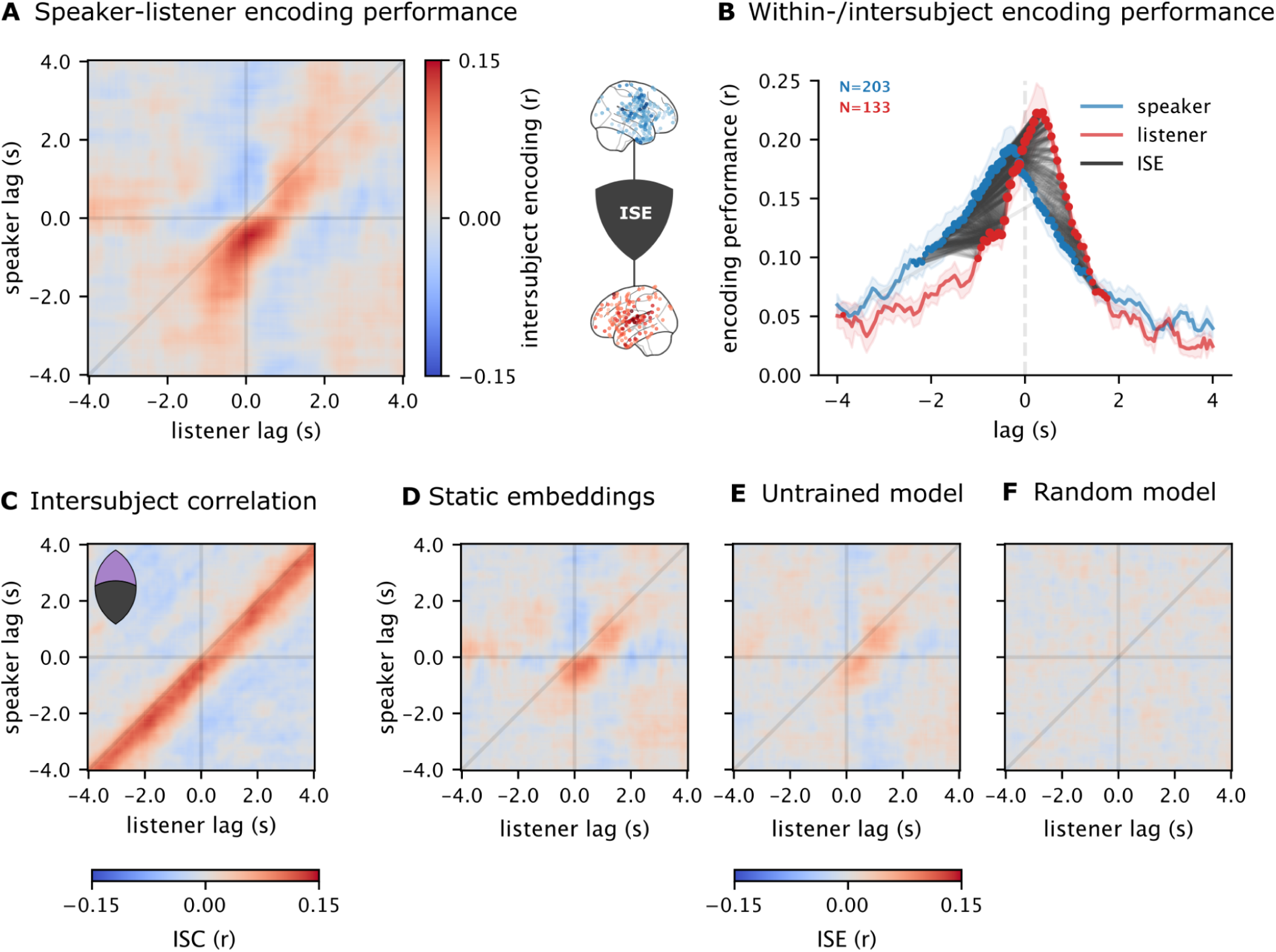
Speaker–listener brain-to-brain linguistic coupling. (**A**) Time-resolved speaker–listener encoding performance across all electrodes and regions for each pair of lags (see Fig. S1 for statistical thresholding). This plot corresponds to the shared variance between the speaker and listener’s brain activity captured by the linguistic encoding model (black intersection in Fig. 1B). Speaker–listener encoding peaks in the bottom right quadrant, indicating that linguistic content emerging in the speaker’s brain prior to word articulation re-emerges in the listener’s brain after word articulation. (**B**) Within-subject encoding performance for the speaker (blue) and listener (red). Gray lines connect significant pairs of lags in the speaker–listener encoding performance matrix (panel A). Line transparency indicates correlation strength, while the size of each circle denotes the overall magnitude for that lag (L2-norm of row or column). (**C**) Speaker–listener intersubject correlation (ISC) without an explicit language model. This analysis directly measures the correlation between word-level neural activity but cannot isolate word-by-word the linguistic content driving brain-to-brain coupling. (**D**) Intersubject encoding performance using non-contextual, static embeddings from the trained LLM. (**E**) Intersubject encoding performance using word embeddings from an untrained version of the same LLM and (**F**) random embeddings with no linguistic structure.

### Speaker–listener linguistic coupling is word-, context-, and conversation-specific

During face-to-face communication, the speaker–listener brain responses can be coupled due to other variables, such as facial expressions, gestures, and background sounds that are not strictly linguistic in nature. We next computed the direct intersubject correlation (ISC) between the speaker and listener’s brain activity (i.e., not mediated through a language model) in a way that matched the folding scheme used for cross-validation in the encoding analysis. Replicating prior results (Stephens et al., 2010; Dikker et al., 2014; Silbert et al., 2014; Liu et al., 2017; Meshulam et al., 2021; Nguyen et al., 2022; Davidesco et al., 2023). We found a strong coupling between speaker and listener neural activity during natural conversations. Direct coupling was consistently below the diagonal, indicating that the speaker’s brain activity preceded the listener’s brain activity (Fig. 3C). ISC analysis cannot isolate the word-by-word linguistic content of the conversation and therefore is not locked to the moment of articulation of each word. Words occurring before and after the current word, regardless of their content, will contribute to the observed correlation, yielding high correlations all along the diagonal. The model-based speaker–listener linguistic encoding results (Fig. 3A), on the other hand, are temporally-specific, suggesting that the embeddings capture more word-specific linguistic coupling than can be observed using ISC analysis.

We next asked whether speaker–listener linguistic coupling was sensitive to the specific meaning of words in context. We extracted non-contextual, static embeddings from GPT-2 with the same dimensionality as the contextual embeddings. In this setting, each occurrence of a given word receives the same embedding, capturing the “average” meaning of that word across all contexts. Similar to other types of word embeddings—such as word2vec and GloVe (Mikolov et al., 2013; Pennington et al., 2014)—these representations cannot capture the unique meaning of words in context. For example, “cold” will receive the same embedding regardless of whether the context refers to the personality trait or temperature. We found that speaker–listener linguistic coupling was significantly stronger for contextual embeddings over non-contextual word embeddings (Fig. 3D; Fig. S3). Furthermore, to demonstrate that our results are not limited to our choice of an autoregressive language model (GPT-2), we replicated our core results using BERT (Devlin et al., 2018), a well-studied masked language model (Fig. S4).

To ensure that the model-based speaker–listener coupling was driven by the linguistic structure of the contextual embedding space, we performed two additional control analyses. In the first analysis, we extracted “embeddings” from an *untrained* GPT-2 model for the same word-by-word input as the trained model used in the main analysis. The encoding analysis with untrained embeddings can capture certain statistical features of the input, such as the repeated co-occurrence of individual words in similar contexts; in addition, the untrained model preserves the positional encoding scheme of the trained model, capturing word-position information. Nonetheless, the untrained model yielded significantly lower intersubject speaker–listener encoding performance (Fig. S3); the model-based predictions accounted for only a small proportion of variance between speaker and listener (Fig. 3E). In the second analysis, we generated random normal vectors (differing across each word instance) with the same dimensionality as the actual embeddings. Random embeddings do not encode meaningful linguistic geometry and yielded near-zero intersubject speaker–listener encoding performance (Fig. 3F).

Finally, we asked whether a given speaker and listener tend to explore a conversation-specific region of the linguistic space. To examine the uniqueness of linguistic coupling in each conversation, we compared the speaker and listener weight matrices estimated by the encoding models within and across conversations. We found that the relationship between speaker and listener model weights is specific to each dyadic conversation and does not generalize across conversations; that is, each conversation was biased toward a particular subset of features in the contextual embedding space (Fig. S5).

### Linguistic coupling across language areas within speakers and listeners

We next used LLM contextual embeddings to assess linguistic coupling across regions of the cortical language network within speakers and listeners. We correlated model-based predictions from one region with the actual neural activity in other regions. For example, we used encoding models trained on neural activity in the speaker’s ATL to predict neural activity in the speaker’s STG. Similarly, we used encoding models trained on neural activity in the listener’s STG to predict neural activity in the listener’s ATL. This analysis yielded lag-by-lag encoding matrices across pairs of language regions within the speaker and the listener (Figs. S6, S7).

During language production, we found a dense network of inter-regional linguistic encoding with many regions encoding similar features of the linguistic embedding space (Fig. 4A, top). These inter-regional connections include both higher-level (ATL, SMG) and lower-level (SM, STG, IFG) language areas. For example, the speaker’s STG and IFG are both coupled to the speaker’s SM but with different temporal structures (Fig. 4A, bottom; Fig. S6); SM precedes STG, and coupling is closely tied to word articulation, while linguistic coupling between SM and IFG is largely synchronous and more temporally diffuse. Interestingly, SM tends to precede most regions, except for IFG. In contrast, during language comprehension, linguistic coupling reveals a more sparse network comprising typical language areas (Fig. 4C): STG is coupled with and generally precedes IFG, MTG, and ATL (Fig. 4C, bottom). Unexpectedly, while SM did have strong encoding performance during comprehension (Fig. 2B), it was not strongly coupled to any other region, suggesting that it encodes a particular set of linguistic features not shared with other areas (Fig. S7).

**Fig. 4.**
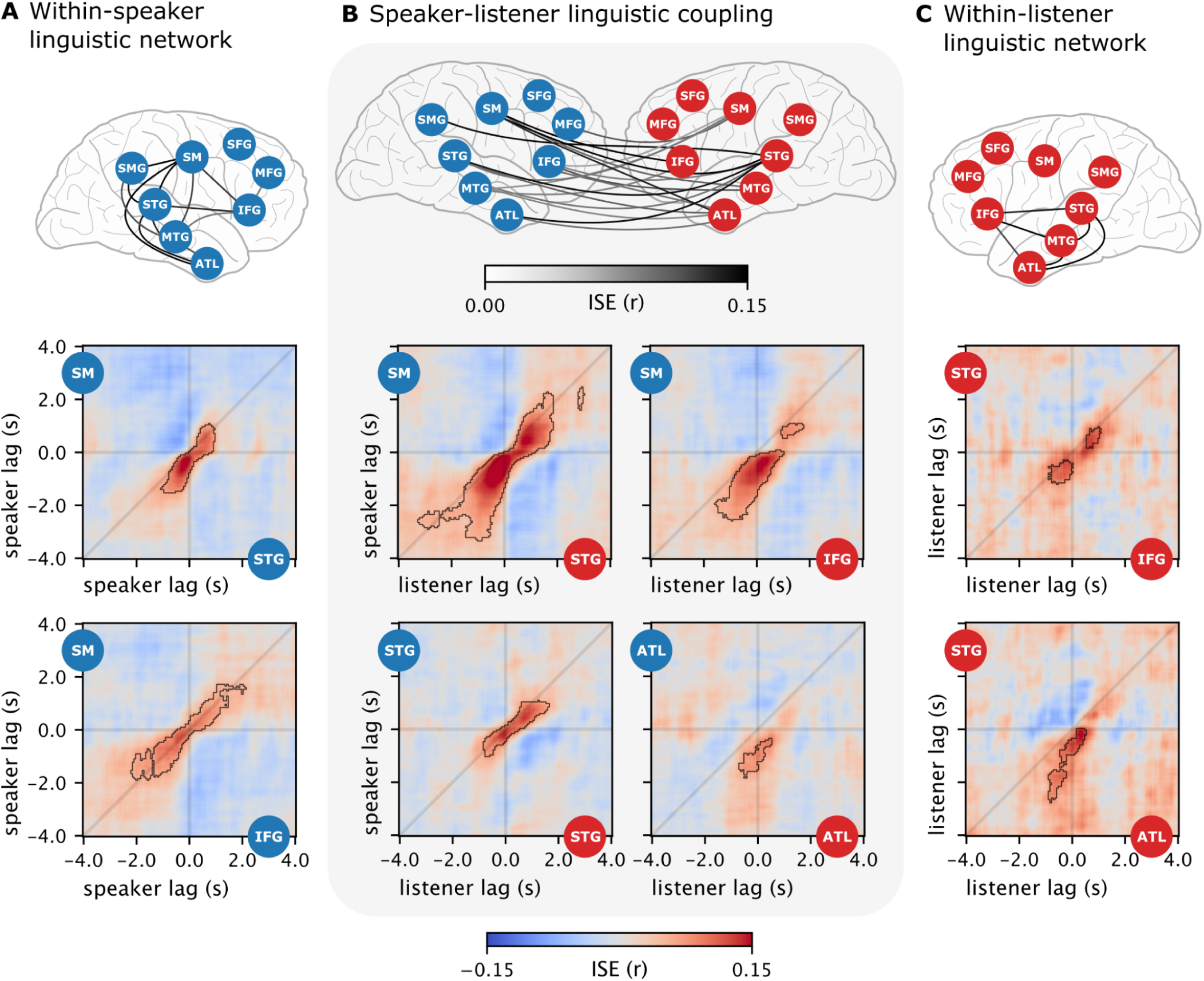
Inter-regional linguistic coupling within and across subjects. (**A**) Linguistic encoding across regions of the cortical language network within the speaker’s brain. The connection between the anterior temporal lobe (ATL) and somatosensory (SM), for example, indicates the maximum encoding performance across lags for a model trained on ATL and tested on SM (and vice versa). The network diagram (top) depicts significant coupling between two regions as bidirectional gray lines (*p* < .05, Bonferroni corrected) where the darkness of the connection indexes the strength of linguistic encoding. Below are lag-by-lag within-speaker encoding matrices for two example pairs of regions with contour lines denoting significant lags (see Fig. S6 for all pairs of ROIs and the number of electrodes per area). (**B**) Intersubject encoding performance between the speaker (blue) and listener (red) across multiple language areas. Network diagrams (top) show significant speaker–listener coupling for pairs of regions; the darkness of the connection indexes the strength of linguistic encoding. Below are speaker–listener lag-by-lag encoding matrices for four example pairs of regions. (**C**) Linguistic encoding across regions of the cortical language network within the listener’s brain (top), with lag-by-lag encoding matrices for two example pairs of regions below.

### Speaker–listener linguistic coupling across language areas

Speaker–listener intersubject linguistic coupling was widespread across the cortical language network and partially asymmetric between different language areas (Fig. 4B). In the speaker, SM, STG, and ATL were the primary drivers of the speaker–listener coupling; in the listener, STG, IFG, and ATL were the primary receivers (Fig. S8). The speaker’s SM was strongly coupled with the listener’s STG, IFG, and ATL. Interestingly, the speaker’s SM activity before word onset was coupled to the listener’s STG before word onset (lower left quadrant), then differently coupled after word onset (upper right quadrant, with minimal coupling in the lower right quadrant); this suggests that both regions encode shared features that change rapidly at word articulation. In contrast, the speaker’s SM before word onset was mostly strongly coupled to the listener’s IFG after word onset (lower right quadrant). While most speaker–listener couplings were asymmetric between language areas, the speaker’s STG was coupled to the listener’s STG along the diagonal around word articulation, suggesting a short delay in these lower-level regions (Fig. 1C). The speaker’s ATL prior to word onset was coupled with the listener’s ATL around word onset with a larger speaker–listener latency (farther off the diagonal). These results suggest a network configuration where linguistic coupling between the high-level language and articulatory areas drives speech production, whereas coupling primarily between STG and IFG, followed by higher-level regions, underlies speech comprehension.

## Discussion

In the current study, we show that contextual embeddings from a large language model (LLM) encode the shared linguistic space between speakers, allowing us to quantify how linguistic information is communicated from brain to brain in spontaneous, face-to-face conversations. Specifically, for each word in the conversation, the linguistic embedding space can be used to recover the word-specific brain activity shared between the speaker and listener. Moreover, we find that linguistic content emerges in the speaker’s brain activity before word articulation, and the same linguistic content later re-emerges in the listener’s brain after word articulation. This temporal dynamic matches the overall flow of information in a conversation, where conversants dynamically alternate roles in transmitting their ideas to one another. LLMs provide a new computational framework for learning the shared linguistic code—i.e., the mutual understanding of the meaning of words in context—that humans use to communicate effectively.

Our findings align with prior work demonstrating that LLM contextual embeddings capture linguistic features of brain activity during language comprehension (Fig. 2, red) (Toneva & Wehbe, 2019; Heilbron et al., 2020; Schrimpf et al., 2021; Goldstein, Zada, et al., 2022; Kumar et al., 2022; Caucheteux & King, 2022; Cai et al., 2023). Here, we demonstrate that these embeddings also capture the emergence of linguistic features—principally before word articulation—during spontaneous speech production (Fig. 2, blue). This suggests that the cortical processes supporting speech production and comprehension tap into a common set of relatively abstract, context-specific linguistic features reflecting the statistical structure of real-world language (in large text corpora) (Pickering & Garrod, 2013). The word-by-word temporal specificity (Fig. 2B) and importance of context (Fig. S3) suggest that each word is rapidly processed, incorporated into the predictive context, and then rapidly integrated with the next word (Hasson et al., 2015; Christiansen & Chater, 2016; Goldstein, Zada, et al., 2022). Unlike prior work, which has largely relied on rehearsed or professionally-produced language stimuli (e.g., podcasts or public recitals by professional storytellers), our results are derived from free-form, interactive dialogues, providing evidence for broader applicability of contextual embeddings for modeling neural activity in naturalistic, conversational environments.

Prior work has demonstrated that the speaker’s and listener’s brains are coupled during verbal communication (Stephens et al., 2010; Jiang et al., 2012; Silbert et al., 2014; Dikker et al., 2014; Liu et al., 2017; Bevilacqua et al., 2019; Meshulam et al., 2021; Nguyen et al., 2022). Our findings replicate this direct brain-to-brain coupling during natural conversations (Fig. 3C), but for the first time, with the spatiotemporal resolution of ECoG recordings in dyadic pairs of epilepsy patients. Direct brain-to-brain coupling, however, is content agnostic: any shared stimuli, including non-linguistic variables that are not related to the conversation, can induce correlated activity across two brains. Moreover, in our dataset, direct intersubject coupling does not capture the temporal specificity of word-by-word linguistic content revealed by the intersubject linguistic encoding analysis (Fig. 3A). In contrast, our novel intersubject speaker–listener encoding framework models word-specific variance in brain-to-brain coupling as mediated by an explicit language model. That said, direct coupling tends to be stronger than intersubject linguistic encoding; this indicates that there are components of brain-to-brain coupling not captured by our linguistic encoding model. Future work may capitalize on our intersubject encoding framework to incorporate models of speech features, intonation, prosody, and visual features like gesture and facial expressions to further account for unexplained variance in brain-to-brain coupling.

Large language models represent and process language in a radically different way than the rule-based, symbolic language models of classical psycholinguistics. Several core ingredients make these models particularly useful for examining the neurocomputational underpinnings of natural language processing in the human brain (Richards et al., 2019; Hasson et al., 2020). First, these models learn from real-world language generated by other humans using straightforward objectives, like predicting forthcoming words, and, more recently, using reinforcement learning based on conversational feedback (Ouyang et al., 2022). These objectives are readily available to all language users and may play a critical role during online speech processing in the human brain (Goldstein, Zada, et al., 2022). Second, like the brain, these models use a population code to represent each word in a high-dimensional embedding space distributed across relatively simple computing elements (Goldstein, Dabush, et al., 2022). Third, the architecture of large language models enables them to express the richness and complexity of real-world language; these models learn to represent the unique meaning of words in context. The present data provide a singular challenge for these models: unrehearsed, free-from conversations where words are spontaneously generated, meaning shifts with ongoing context, and nothing is strictly repeated (Nastase et al., 2020). Nonetheless, these models successfully capture context-specific, word-by-word linguistic content encoded in the brain activity of both speaker and listener. Our findings suggest that these three ingredients may allow these models to tap into the same shared linguistic “code” that humans use to communicate our thoughts to one another.

## Materials and Methods

### Electrocorticography acquisition

Twelve participants (6 dyads) engaged in free-form, first-time conversations (Table S2). One dyad was excluded from the analysis due to a short conversation length (3.56 minutes) and an insufficient number of words spoken (185 and 156 words for each participant, respectively). Participants were instructed to discuss any topic, including hobbies, vacation stories, movies, etc. Participants were recruited from the New York University School of Medicine Comprehensive Epilepsy Center and provided oral and written informed consent before study participation, according to the New York University Langone Medical Center Institutional Review Board. All participants elected to undergo intracranial monitoring for clinical purposes and were informed that their clinical care was unrelated to participation in this study and that withdrawing from the study at any point would not affect their medical treatment.

Electrode placement was determined by clinicians for each participant based on clinical criteria. Brain activity was recorded from intracranially implanted subdural platinum–iridium electrodes embedded in silastic sheets (2.3-mm-diameter contacts; Ad-Tech Medical Instrument). Decisions related to electrode placement and invasive monitoring duration were determined solely on clinical grounds without reference to this or any other research study. Electrodes were arranged as grid arrays (8 × 8 contacts, 10 mm center-to-center spacing) or linear strips.

Recordings from grid, strip, and depth electrode arrays were acquired using the NicoletOne C64 clinical amplifier (Natus Neurologics), band-pass filtered from 0.16–250 Hz, and digitized at 512 Hz. Intracranial electroencephalography signals were referenced to a two-contact subdural strip facing toward the skull near the craniotomy site. All electrodes were visually inspected, and those with excessive noise artifacts, epileptiform activity, or no signal were removed from subsequent analyses.

Presurgical and postsurgical T1-weighted magnetic resonance imaging (MRI) scans were acquired for each participant, and the location of the electrodes relative to the cortical surface was determined from co-registered magnetic resonance imaging or computed tomography scans (Yang et al., 2012). Co-registered, skull-stripped T1 images were nonlinearly registered to an MNI152 template, and electrode locations were extracted in Montreal Neurological Institute space (projected to the cortical surface) using the co-registered image.

### Electrode localization

Electrodes were localized to anatomically-defined cortical regions based on the Desikan-Killiany atlas in FreeSurfer (Desikan et al., 2006). Regions of interest (ROIs) consisted of one or more of the following atlas labels: somatomotor (SM) cortex, including precentral and postcentral gyri; superior temporal gyrus (STG), including the posterior superior temporal sulcus; middle temporal gyrus (MTG); middle frontal gyrus (MFG), comprising rostral and caudal middle frontal gyrus; supramarginal gyrus (SMG); inferior frontal gyrus (IFG), comprising pars opercularis, pars orbitalis, and pars triangularis; anterior temporal lobe (ATL), comprising anterior inferior temporal cortex and the temporal pole; superior frontal gyrus (SFG). Electrodes overlapping multiple regions were assigned based on their percent overlap with the largest area. Only electrodes in the left hemisphere were considered.

### Signal preprocessing

ECoG data were standardized according to the Brain Imaging Data Structure (iEEG-BIDS) (Holdgraf et al., 2019) and preprocessed using the MNE Python library (Gramfort, 2013), with accompanying customized Python scripts. The pipeline consisted of the following steps: removing spikes, re-referencing, notch filtering, then high-gamma band extraction. First, large spikes exceeding four quartiles above and below the median of each channel were removed, and replacement samples were imputed using the SciPy *pchip_interpolate* function (Virtanen et al., 2020). We then re-referenced all electrodes in each subject to account for shared signals across channels using an independent component analysis method (Michelmann et al., 2018). We removed line noise at 60, 120, and 180 Hz frequencies using MNE’s *notch_filter* function with a width of 2 Hz. Finally, we estimated broadband power using an IIR filter followed by a Hilbert envelope computation for the 70-200 Hz frequency band (using *filter* and *apply_hilbert(envelope=True)* from MNE). High-frequency broadband power has been shown to correlate with local neural firing rates (Jia et al., 2013; Muller et al., 2016).

### Transcription and alignment

We recorded each conversation’s audio with a microphone at the full 44,100 Hz rate. In addition, the clinical amplifier also received the microphone output and was saved at 512 Hz. Each conversation was manually transcribed, and each utterance was manually identified to a speaker and aligned to the audio. Sounds such as laughter, breathing, or inaudible speech were marked to improve the alignment’s accuracy. Punctuation and capitalization were included to the best ability of the transcriber. The Montreal Forced Aligner (McAuliffe et al., 2017) was used to align the audio to the transcript automatically and to compute onsets and offsets for each word. The number of words per conversation is reported in Table S3. To finely align the audio with the brain activity, we cross-correlated the microphone audio (44,100 Hz) envelope with the clinical amplifier audio (512 Hz) envelope. The lag of the peak correlation was used to translate the word onsets to match the brain data. We further validated this alignment in two ways: first, by cross-correlating the ECoG signal with a downsampled version of the audio envelope. Most electrodes in STG exhibited clear peaks with small latency in the correlogram. Second, we extracted epochs around the onset for each word and averaged them into one “evoked” response. This showed a clear peak after onset in the audio envelope and in several electrodes.

### Contextual embedding extraction

We used the pre-trained extra-large version of GPT-2 (Radford et al., 2019) with 48 layers in the HuggingFace Transformers library (Wolf et al., 2020). We first converted all words from the raw transcript to GPT-2-specific tokens (full words and subwords) and then to integer identifiers. We supplied the model with each token up to the maximum allowed by the 1024-token context window to extract the embedding from the activations (hidden states) of the final word in the sequence. Embeddings at each layer are 1,600-element numerical vectors. Only the middle 24th layer was considered in subsequent analyses, as middle layers have been shown to be better predictors of neural activity (Caucheteux & King, 2022; Goldstein, Ham, et al., 2022). Finally, sub-word token embeddings were averaged for each whole word to harmonize with the original transcript. To compare embedding spaces, this same procedure was used to extract embeddings from an untrained GPT-2 model—with random initial weights but the same architecture, context window, and inputs. We also extracted static, non-contextual GPT-2 embeddings corresponding to the token embedding weights learned by the model. As a final control, we generated random normal embeddings with the same dimensionality as GPT-2 embeddings (1,600 features) for each word instance: thus, two instances of the same word would receive two separate random embeddings (to mimic the fact that actual GPT-2 embeddings differ for different instances of the same word across contexts).

In order to demonstrate that our results generalize to other large language models that also capture a similar shared linguistic space, we replicated our core analyses using the widely studied masked language model BERT (Devlin et al., 2018). We used the large, cased, whole-word-masking version of BERT from the HuggingFace library. We divided each transcript into utterances composed of one or more sentences. One speaker produced each utterance at a time. Then, each utterance was prepended with the words that appeared before it, if any, to fill up the maximum context length of the model (512 tokens). Once fed into the model, we extracted the activations corresponding only to the utterance in consideration of that input and not the context. These activations were taken from the middle layer only to match our procedure with GPT-2.

### Encoding analysis

A linear model was estimated using ridge regression to predict the neural signal separately for each electrode and every lag relative to word onset. We used the full 1,600 contextual embedding from the language model as the predictors. Two encoding models were estimated for each subject: one for words they produced as a speaker and a separate model for words they heard as a listener. The neural signal was divided into epochs for every word (separately for spoken or heard words). Each epoch ranges from –4 s to +4 s in 129 bins of 250 ms overlapping frames with jumps of 62.5 ms (see Fig. S9 for the relation of the 250 ms window size to results). We fit separate encoding models per lag to resolve the temporal dynamics of linguistic encoding. The encoding model was evaluated by computing the Pearson correlation coefficient between actual and predicted neural signals for left-out test sets using 10-fold cross-validation. The data was split consecutively (i.e., into temporally contiguous segments) for cross-validation so as to minimize the autocorrelation between training and test folds. The final correlation values reported in the text are the average correlation across all test folds. We used the *RidgeCV* implementation from the *Himalaya* Python library (la Tour et al., 2022) to fit the encoding models. Regularization coefficients (i.e., L2 penalty terms) were selected from 20 log-spaced values ranging from 1 to 1,000,000 using random search with nested cross-validation (5-fold inner cross-validation for hyperparameter selection within each training fold of the outer 10-fold cross-validation loop; as implemented in *Himalaya*). Note that for both the predictor matrix of embeddings and the target vector for each electrode, the number of samples corresponds to the number of words (not the number of time points).

### Electrode selection

We first performed a permutation test to generate a null distribution based on phase-randomized neural signals to select electrodes involved in language production and comprehension for further analysis. Phase randomization effectively decouples the time series of neural activity from the onsets/offsets of each word and utterance. For each of the 1,000 permutations, we fit the same ridge-regression encoding models described above based on static, non-contextual GPT-2 embeddings across all electrodes and lags (Fig. S10). The null distribution was constructed from the maximum encoding performance across lags; one *p*-value per electrode was calculated according to its own null distribution. Constructing the null distribution from maximum encoding performance values across lags effectively controls for multiple tests across lags (Nichols & Holmes, 2002). We then controlled the false discovery rate (FDR; Benjamini & Hochberg, 1995) at an alpha value of 1% to control for multiple tests across the *p*-values at each electrode for production and comprehension in each subject. The final number of selected electrodes per subject per region is detailed in Table S4. Of all selected electrodes, 231 were from grids, 87 were from strips, and 18 were depth electrodes.

### Intersubject encoding

We used the trained encoding models (i.e., the learned weight matrices) from the within-speaker and within-listener analyses to assess intersubject encoding. Specifically, we used the predicted neural signal from a model trained on speaker data (the production model) and correlated it with the actual neural signal of the listener (Toneva et al., 2022). Because the production models are estimated separately at each lag relative to word onset (and therefore yield different predictions for each lag), we computed correlations between the model-predicted activity and actual neural activity at each pair of lags ranging from –4 s to +4 s between speaker and listener. This analysis is not strictly symmetric if performed in the opposite direction: model-based predictions from the encoding model estimated in the listener (the comprehension model) evaluated against the speaker’s actual neural activity. For this reason, we performed the same intersubject encoding analysis for the listener: we computed comprehension model predictions derived from the listener, correlated these with the speaker’s actual neural activity, then averaged these correlations with those computed from the production model and evaluated against the listener’s neural activity. In practice, the results for both directions are very similar (Fig. S1).

This analysis yields model-based predictions (and the corresponding actual neural activity) for every word at each lag and each electrode. To construct a lag-by-lag matrix that summarizes across electrodes (e.g., across all electrodes as in Fig. 3, or across electrodes within a given language area as in Fig. 4), we averaged the model-predicted activity and actual activity (retaining lags) prior to computing the correlation between predicted and actual neural activity. This correlation value is visualized at each pair of lags. We perform this process for each speaker–listener pair, and the final correlations are summarized across dyads using a weighted average, where the weights correspond to the relative number of words for each dyad; this weighted averaging procedure mitigates variance in correlations derived from small training/test folds in dyads with fewer words. Each row of the resulting matrix indicates how well the model-based predictions for a given speaker lag (the row index) match the listener’s brain activity at each lag. The diagonal represents the same (i.e., matching) lag between speaker and listener; anything below the diagonal indicates that the speaker precedes the listener. The bottom half of the matrix corresponds to speaker–listener linguistic coupling based on the speaker’s brain activity prior to word onset. The right half of the matrix corresponds to the speaker–listener linguistic coupling based on the listener’s brain activity after word onset. The bottom right quadrant of the matrix corresponds to pairs of lags where the linguistic content of the speaker’s brain prior to word onset is coupled to the linguistic content of the listener’s brain after word onset.

This same procedure was applied across regions within-speaker and within-listener. Usually, encoding models within subjects are trained on one electrode (and one lag), and the correlation between the model’s predictions and the actual activity of the same electrode (and lag) is evaluated on a held-out test set. Following the same procedure as above, we can evaluate the encoding model predictions for one region against another region. This tests whether two brain regions in the speaker (or listener) use the same linguistic features from the embedding space. We ran this procedure for all pairs of regions within-speaker (Fig. 4A; Fig. S6) and within-listener (Fig. 4C; Fig. S7).

For comparison, we computed intersubject correlation (ISC; Simony et al., 2016; Nastase et al., 2019) by applying the same procedure as above, but instead of using predicted neural activity from encoding models, we computed the correlations between the actual speaker neural activity and actual listener neural responses for the corresponding test sets.

### Statistical significance

To evaluate the statistical significance of the lag-by-lag intersubject encoding matrix, we generated a null distribution of intersubject encoding for each pair of lags based on phase-randomized neural signals. We re-estimated encoding models for each of the 1,000 phase permutations. For the whole-brain analysis (Fig. 3A), we computed 1,000 iterations of the lag-by-lag correlation matrix using the phase-randomized models for both speaker and the listener. Specifically, we correlated the speaker’s phase-randomized model’s predicted neural activity with the listener’s *actual* neural activity and vice-versa (i.e., the same as intersubject encoding, except with the perturbed model). This procedure used the same selected electrodes as the non-randomized analyses. The maximum value across all pairs of lags in each matrix was submitted to the null distribution to control for multiple tests across pairs of lags. Lag pairs were considered significant at an alpha value of 1% based on this distribution, corresponding to an intersubject encoding performance (correlation) value of .052 (*p* < .01, FDR corrected) (Fig. 3B; Fig. S2).

To evaluate the significance of inter-regional intersubject encoding analysis (Fig. 4), we generated a null distribution using the encoding models estimated from phase-randomized neural data. The lag-by-lag intersubject encoding correlation matrix was computed once for each pair of speaker–listener regions (8 x 8), resulting in 1,000 permutation correlations for each pair of regions and each pair of lags (129 x 129). Because of unique electrode coverage per participant and the electrode selection process, the number of subjects and electrodes in each pair of regions differed. To ensure the reliability of these correlations across subjects, we only considered pairs of regions with at least four subjects (Figs. S6, S7, S8). We computed *p*-values for each pair of regions and each pair of lags based on the corresponding null distribution, then used Bonferroni correction to control the family-wise error rate at 5% across pairs of ROIs and pairs of lags. Any resulting pair of ROIs with at least 20 adjacent pairs of significant lags were deemed significant overall (gray lines in Fig. 4). The same statistical analysis was applied to the within-speaker (Fig. S6) and within-listener (Fig. S7) region pairs.

We also tested for a significant difference between intersubject encoding using contextual, model-trained embeddings against static and untrained model embeddings (Fig. 3, Fig. S3). To do this, we compared the observed difference against the 10,000 permutation differences, where we randomly assigned the observation per speaker (10) and per fold (10) to either sample. The resulting *p*-values were correlated for multiple comparisons with FDR at *p* < 0.01.

## Acknowledgments

We thank Bobbi Aubrey, Mariya Toneva, Robert Hawkins, and Diana Tamir for helpful feedback on the data and analyses.

## Author contributions

Conceptualization: ZZ, SAN, AG, UH

Data curation: ZZ, EB, WD, DF, PD, LM, SD, AF, OD

Formal analysis: ZZ

Funding acquisition: UH

Investigation: ES, AP, AZ, LH

Methodology: ZZ, SAN, AYG, SM, UH

Project administration: ZZ, SAN, UH

Software: ZZ

Supervision: SAN, UH

Visualization: ZZ, SAN, UH

Writing – original draft: ZZ, SAN, UH

Writing – review & editing: ZZ, AF, SAN, UH

## Data and materials availability

Code is available at https://github.com/hassonlab/b2b-linguistic-coupling. Due to the sensitive nature of the unconstrained speech, the data cannot be shared publicly; we will make the data available to reviewers upon request.

## Supplementary Materials

**Fig. S1.**
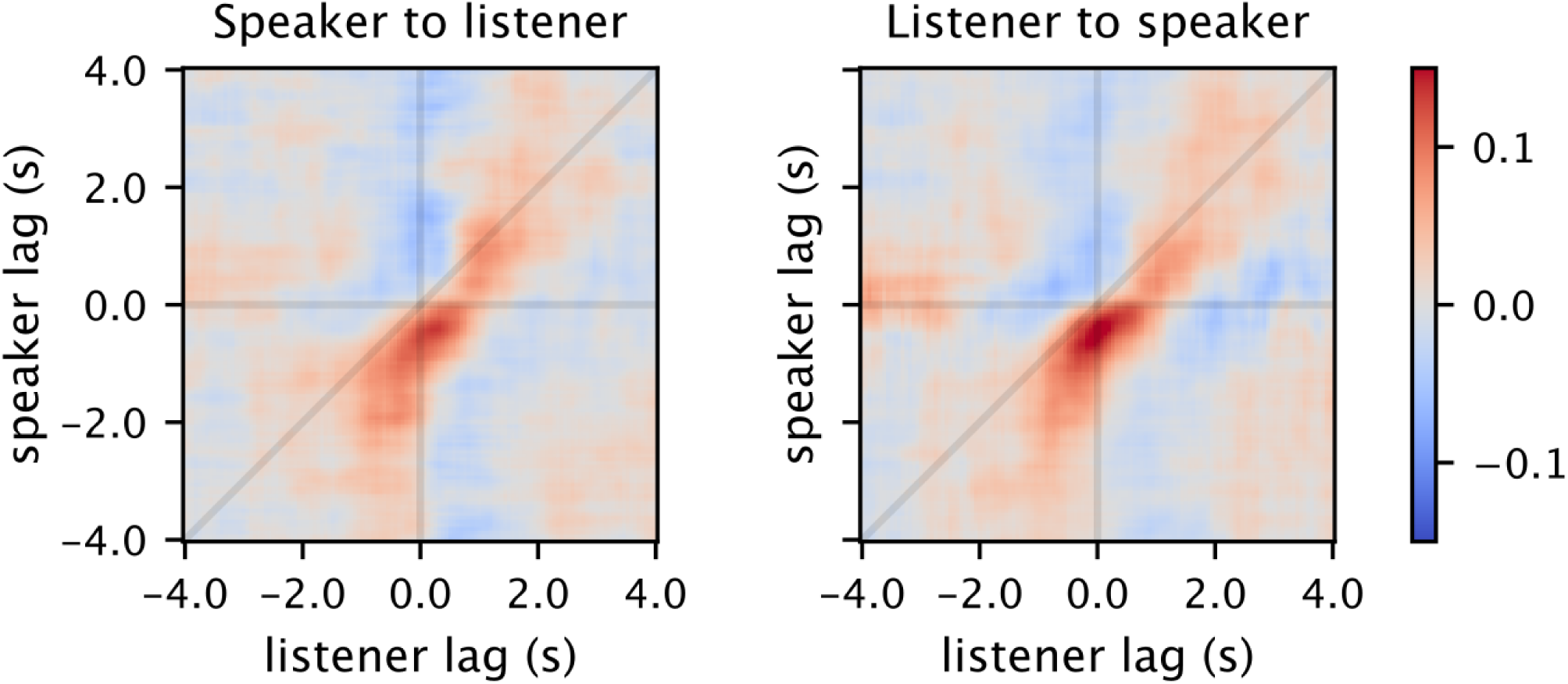
Asymmetric invariance of speaker–listener intersubject encoding. Intersubject encoding performance for models trained on the speaker and tested on the listener (left); intersubject encoding performance for models trained on the listener and tested on the speaker (right). These two “directions” of the encoding analysis correspond to the diagonal black lines from Fig. 1A. This finding suggests that the two directions are qualitatively symmetric; therefore, in the main analyses, we average encoding performance across the two directions.

**Fig. S2.**
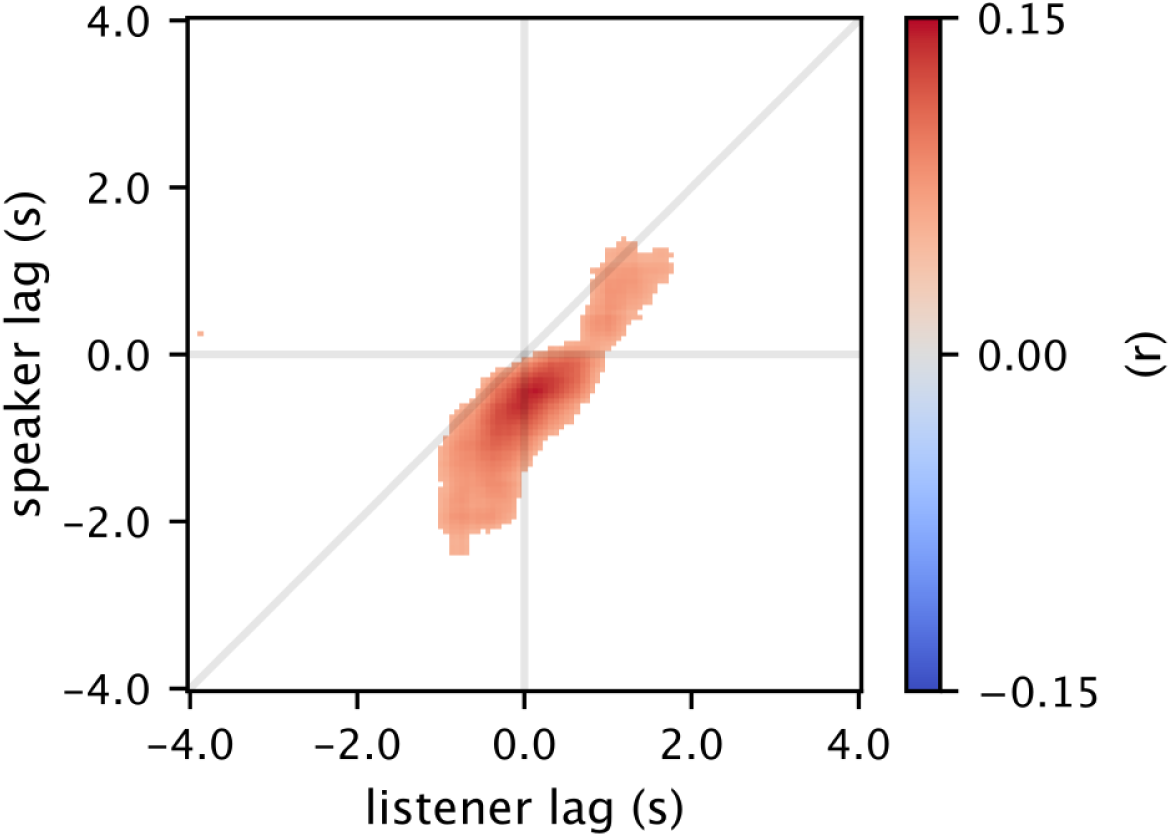
Significant speaker–listener brain-to-brain linguistic coupling. Intersubject encoding performance (correlation) values were thresholded for statistical significance based on a phase-randomization permutation test (*p* < .01, *r* > 0.052). Phase-randomization effectively decouples the time series from the word onsets. The permutation distribution was populated with the maximum encoding performance value across all lag pairs at each permutation iteration, effectively controlling the familywise error rate across all lags.

**Fig. S3.**
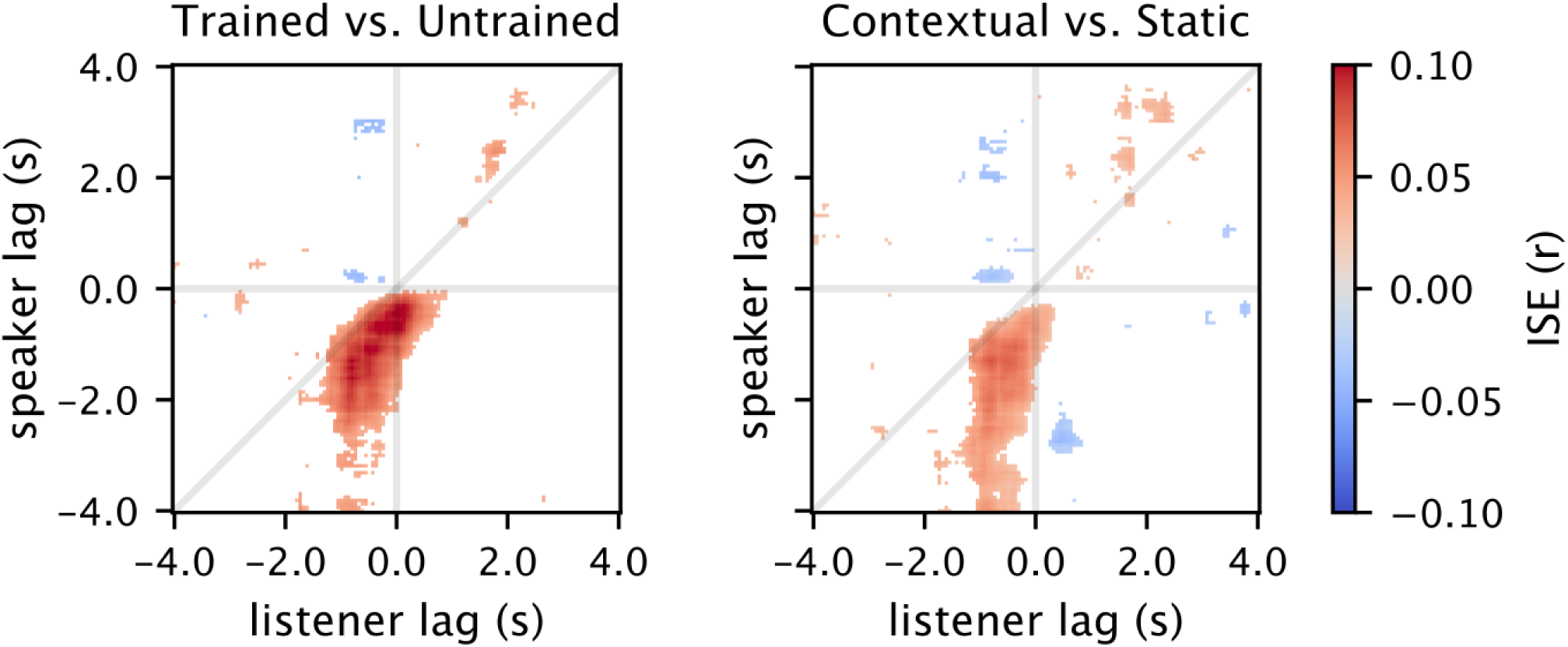
Significant differences in intersubject encoding relative to two control models. The intersubject encoding performance values for the GPT-2 embeddings used in the main analysis were significantly greater than for embeddings extracted from an untrained GPT-model (left). The contextual GPT-2 embeddings used in the main analysis also yielded significantly higher encoding performance than non-contextual, static embeddings extracted from the same GPT-2 model. We ran 10,000 paired samples permutations of the embedding type across subjects and folds for each of the 129 ⨉ 129 lag pairs. Thresholds are based on two-sided *p*-values (*p* < .01, FDR controlled across all pairs of lags).

**Fig. S4.**
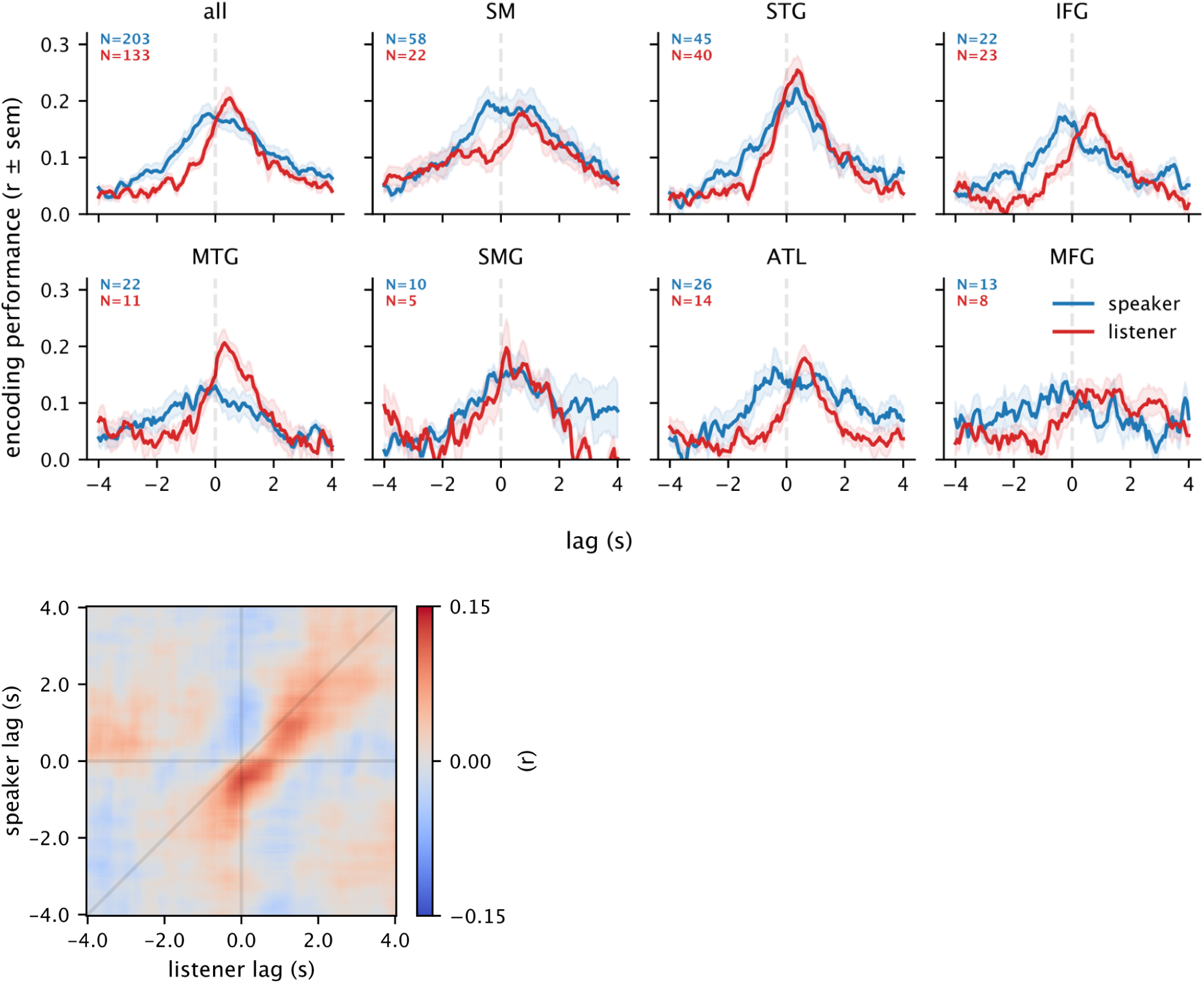
BERT within-subject and intersubject speaker–listener encoding performance. To ensure that our results are not specific to the particular GPT-2 model used in the main analysis, we replicated our main results with word embeddings extracted from BERT (large, 12^th^ layer), a widely-studied, bidirectional, and encoder-based, masked language model. These results suggest that both models similarly capture the shared linguistic space humans use to communicate.

**Fig. S5.**
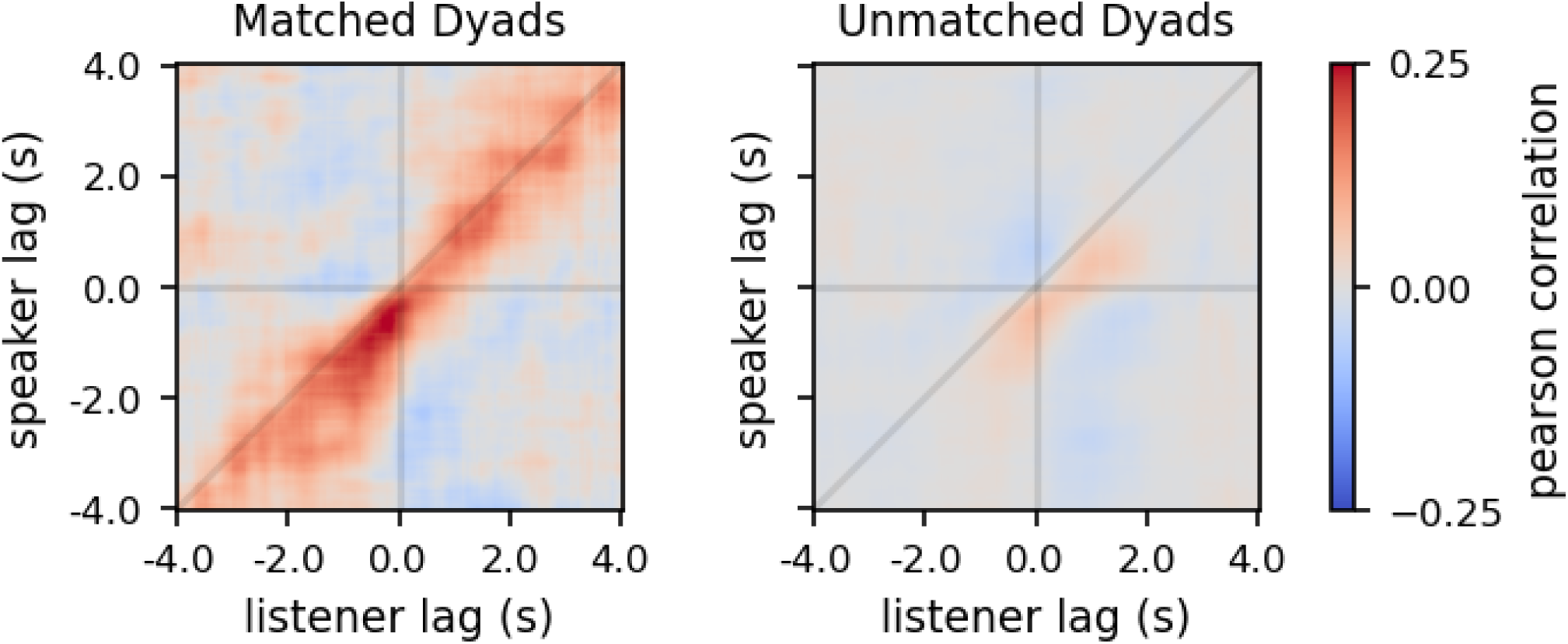
Conversation-specific encoding model weights. To evaluate to what extent each conversation explored a specific region of the linguistic embedding space, we examined the weight matrices learned by the encoding models for each conversation. Correlation of speaker encoding model weights with listener encoding model weights within matched dyads (left) qualitatively exceeded speaker–listener weight matrix correlations between unmatched dyads (right).

**Fig. S6.**
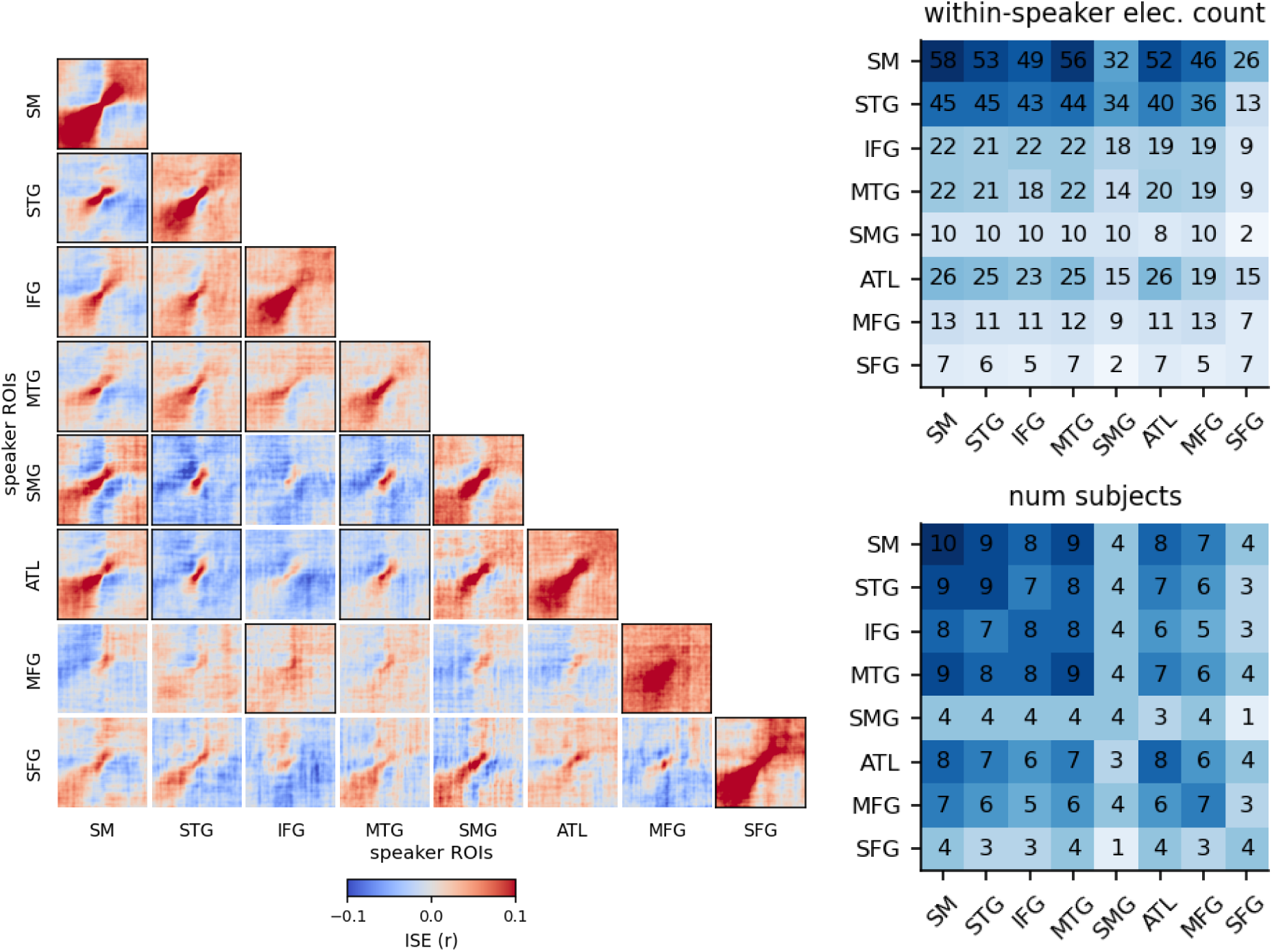
Inter-regional within-speaker encoding. Intersubject encoding performance for each pair of regions within each speaker (left). For example, STG (row) – SM (column) evaluates the model predictions from the speaker’s STG encoding model against the actual activity in the speaker’s SM. Heatmaps with a black border are considered significant (see “Statistical significance” in “Materials and Methods”). At right, matrices indicate the number of electrodes and number of subjects for each pair of ROIs included in this analysis.

**Fig. S7.**
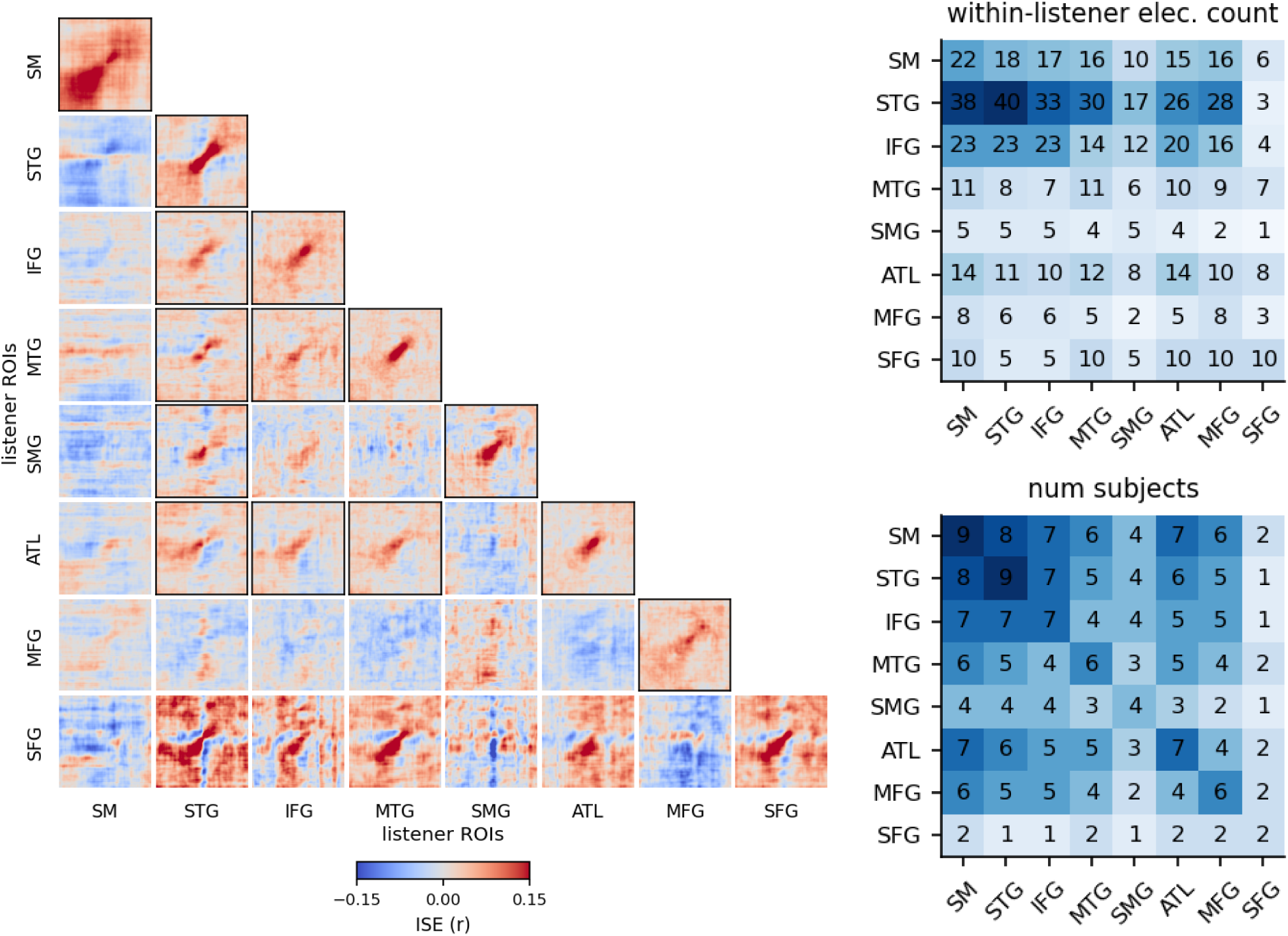
Inter-regional within-listener encoding. Intersubject encoding performance for each pair of regions within each listener (left). Heatmaps with a black border are considered significant (see “Statistical significance” in “Materials and Methods”). At right, matrices indicate the number of electrodes and number of subjects for each pair of ROIs included in this analysis.

**Fig. S8.**
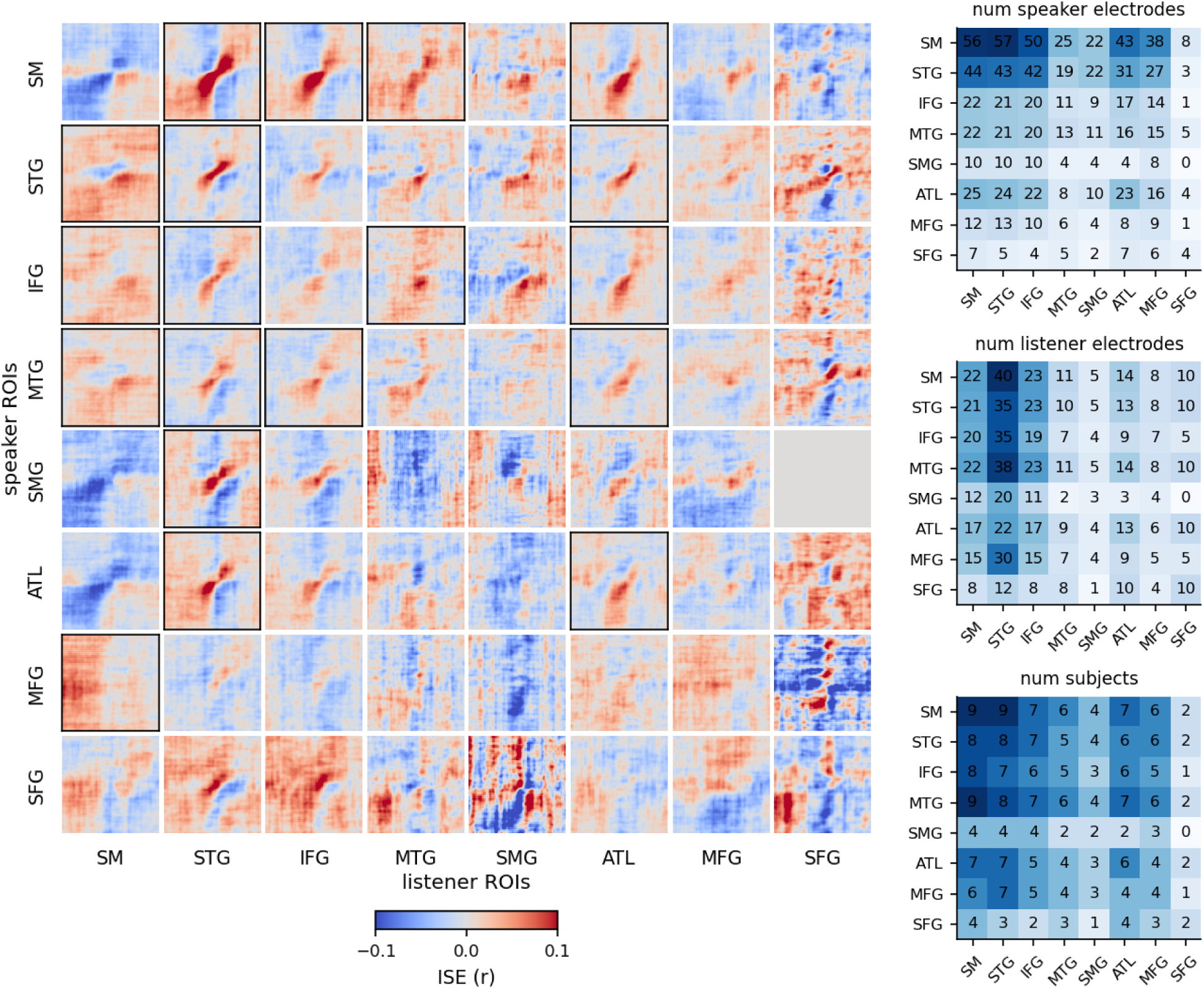
Inter-regional speaker–listener encoding. Intersubject encoding performance for each pair of regions of interest (left). Heatmaps with a black border are considered significant. Due to differences in electrode coverage and selected electrodes, each pair of regions is derived from different pairs of subjects and electrodes. Blue matrices (right) show for each pair of regions the number of speaker electrodes, listener electrodes, and subjects that contributed to the final result. For example, speaker STG to listener MTG has 4 subjects, meaning that 4 ISE heatmaps were averaged for the final result.

**Fig. S9.**
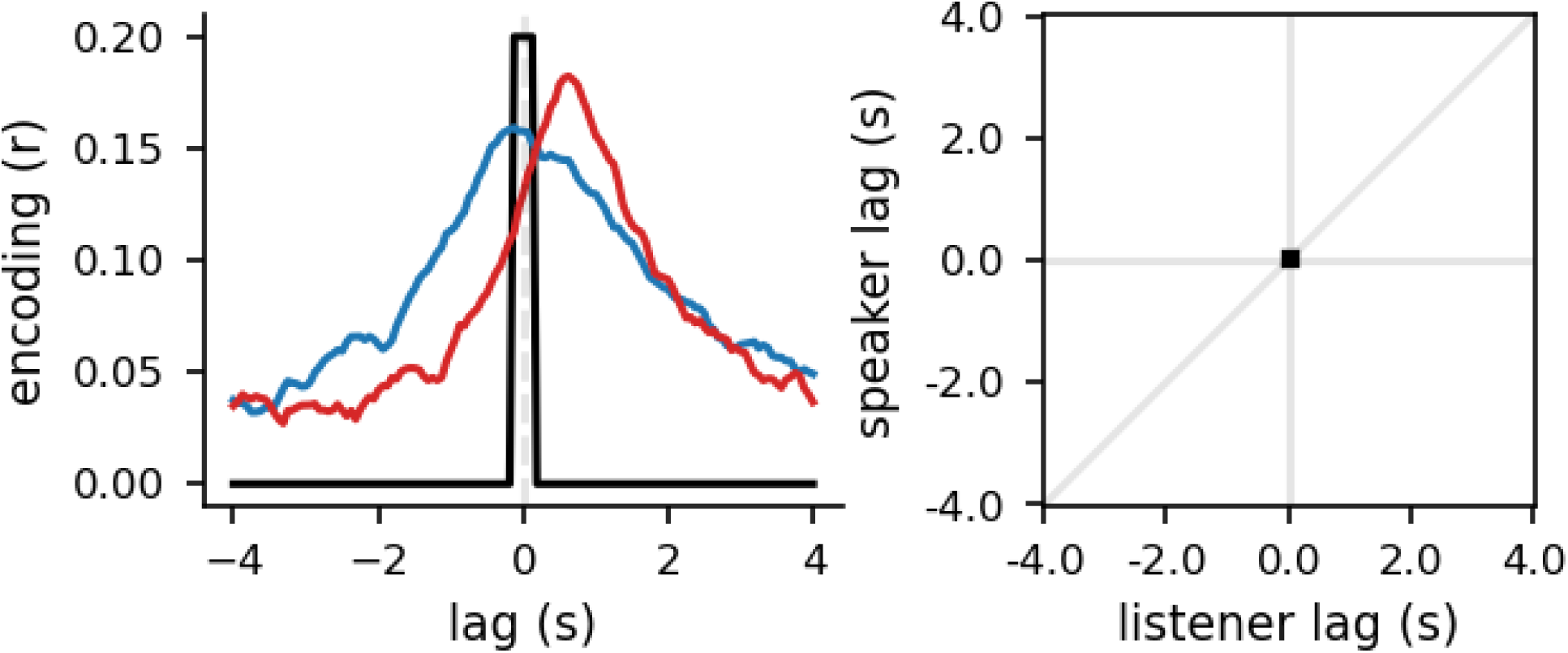
Temporal resolution based on epoch width. The epoching procedure uses overlapping bins, each using a window size of 250 ms to downsample and smooth the signal. The diagram on the left shows the temporal precision of our analysis by epoching an impulse signal for one word at lag 0. The points with signal from the impulse at lag 0 are (-250, 0), (-187.5, 62.5), (-125, 125), (-62.5, 187.5), and (0, 250) milliseconds. On the right is the outer product of the epoch window on the left, showing the temporal precision in the same visualization and axis limits as those in Fig. 3.

**Fig. S10.**
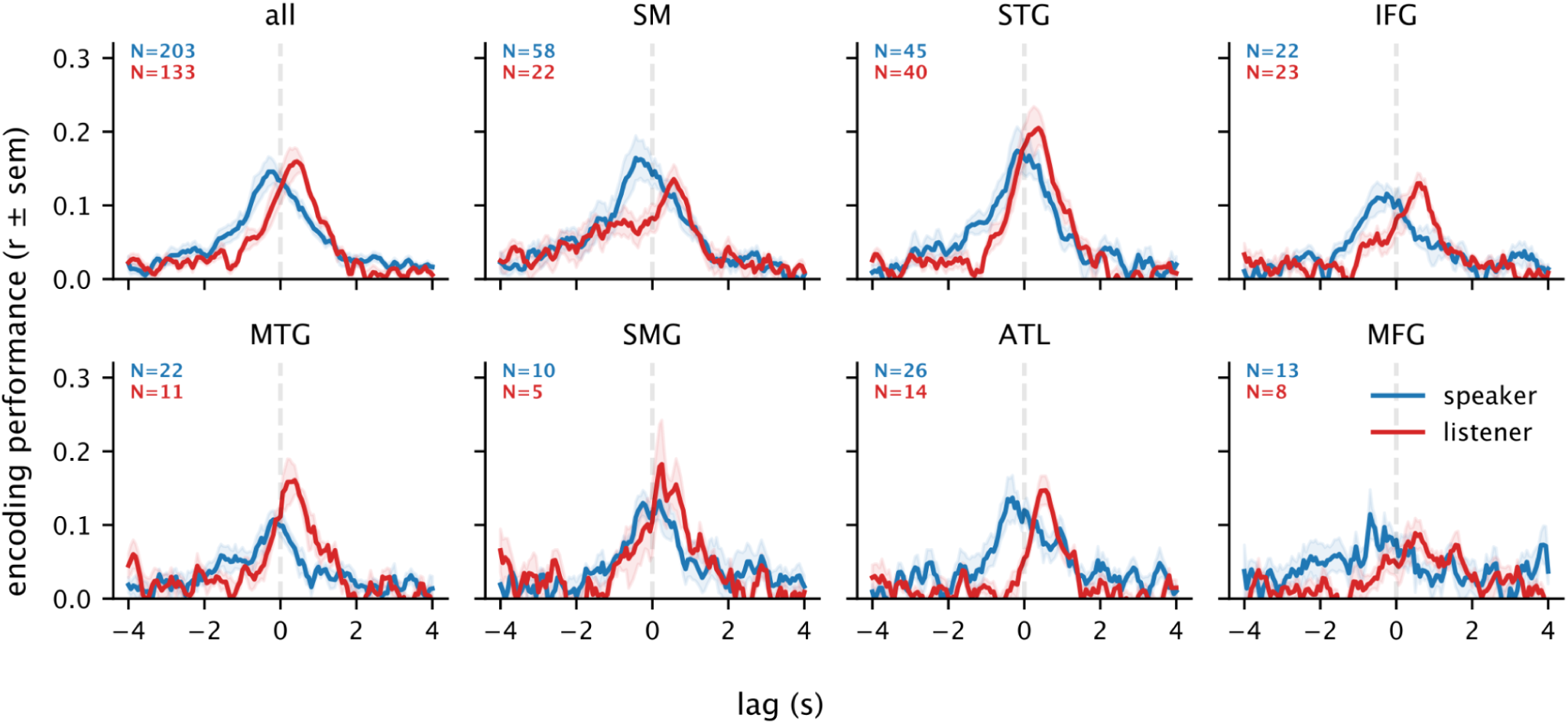
Within-speaker and within-listener encoding performance for static GPT-2 embeddings. For comparison, we replicated the results of Fig. 2B using static embeddings extracted from the token-embedding weights of GPT-2. These non-contextual embeddings yield modest encoding performance.

**Table S1.** Differences in within-subject encoding performance for production and comprehension. We assessed both differences in the temporal lag of peak encoding (relative to word onset) and differences in the magnitude of the peak encoding performance. Statistical significance was assessed based on permutation tests (n = 10,000; permuting assignment of production vs. comprehension for each electrode) for the difference between production and comprehension. The permutation distribution was populated with the maximum peak within-subject encoding performance to control the familywise error rate across lags. For each permutation, each electrode was randomly assigned to either production or comprehension groups, the samples were averaged, and then the maximum correlation and lag differences were retained as the test statistic. Note that each region contained a different number of electrodes. All *p*-values are FDR corrected across ROIs; bolded *p*-values are significant at *p* < .05. Abbreviations: “prod”: production, “comp”: comprehension, “corr”: correlation.

| ROI | prod<br>peak lag<br>(s) | comp<br>peak lag<br>(s) | lag<br>difference<br>(s) | lag<br>difference<br>*p*-value | prod<br>peak corr.<br>(r) | comp<br>peak corr.<br>(r) | corr.<br>difference<br>e (r) | corr.<br>difference<br>*p*-value |
| --- | --- | --- | --- | --- | --- | --- | --- | --- |
| all | -0.2500 | 0.3750 | 0.6250 | **0.00495** | 0.1732 | 0.2158 | -0.0426 | **0.0009** |
| SM | -0.4375 | -1.0000 | 0.5625 | **0.0457** | 0.2004 | 0.1610 | 0.0393 | 0.1473 |
| STG | -0.1875 | 0.2500 | 0.4375 | **0.0272** | 0.1943 | 0.2841 | -0.0898 | **0.0009** |
| IFG | -0.6875 | 0.3750 | 1.0625 | **0.0250** | 0.1549 | 0.1933 | -0.0385 | 0.0648 |
| MTG | -0.5000 | 0.1875 | 0.6875 | 0.0710 | 0.1515 | 0.2305 | -0.0790 | 0.0582 |
| SMG | -0.2500 | 0.1875 | 0.4375 | 0.5080 | 0.1692 | 0.2058 | -0.0366 | 0.4400 |
| ATL | -0.1875 | 0.5000 | 0.6875 | 0.2281 | 0.1745 | 0.2118 | -0.0373 | 0.2310 |
| MFG | -0.3125 | 0.6250 | 0.9375 | 0.1471 | 0.1459 | 0.1649 | -0.0190 | 0.6160 |
| SFG | -1.0000 | 0.3750 | 1.3750 | **0.0050** | 0.1722 | 0.2542 | -0.0820 | 0.0777 |

**Table S2.** Subject demographics and Wechsler Adult Intelligence Scale scores. Scales comprise Verbal Comprehension Index (VCI), Perceptual Organization Index (POI), Processing Speed Index (PSI), Working Memory Index (WMI).

| conversation | subject | sex | age | hand | VCI | POI | PSI | WMI |
| --- | --- | --- | --- | --- | --- | --- | --- | --- |
| 1 | 1 | F | 23 | R | 108 | 107 | 84 | 89 |
|  | 2 | M | 22 | n/a | n/a | n/a | n/a | n/a |
| 2 | 3 | F | 33 | R | 76 | 94 | 71 | 80 |
|  | 4 | M | 49 | R | 121 | 112 | n/a | n/a |
| 3 | 5 | M | 21 | R | 91 | n/a | 86 | 80 |
|  | 6 | F | 58 | R | n/a | n/a | n/a | n/a |
| 4 | 9 | F | 25 | L | 93 | 94 | 97 | 89 |
|  | 10 | F | 20 | L | 112 | 98 | 92 | 105 |
| 5 | 11 | M | 26 | R | 89 | 75 | 65 | 74 |
|  | 12 | F | 22 | L | 74 | 71 | 76 | 69 |

**Table S3.** Conversation statistics per subject. Number of words spoken (production) and number of words heard (comprehension) for each subject in each conversation pair.

| conversation | subject | number of<br>words spoken | number of<br>words heard |
| --- | --- | --- | --- |
| 1 | 1 | 1071 | 2417 |
|  | 2 | 2417 | 1071 |
| 2 | 3 | 1554 | 1341 |
|  | 4 | 1341 | 1554 |
| 3 | 5 | 303 | 2546 |
|  | 6 | 2546 | 303 |
| 4 | 9 | 2205 | 1018 |
|  | 10 | 1018 | 2205 |
| 5 | 11 | 4486 | 1159 |
|  | 12 | 1159 | 4486 |

**Table S4.** Significant electrodes per subject per ROI group. Abbreviations: comp: comprehension (listening); prod: production (speaking).

|  | ROI: | ATL | IFG | MFG | MTG | SFG | SM | SMG | STG |  |
| --- | --- | --- | --- | --- | --- | --- | --- | --- | --- | --- |
| subject | role |  |  |  |  |  |  |  |  | total |
| 1 | prod | 1 | 1 | 2 | 1 | 1 | 5 | 0 | 0 | 11 |
| 2 | prod | 2 | 3 | 2 | 5 | 0 | 11 | 4 | 9 | 36 |
| 3 | prod | 0 | 1 | 0 | 1 | 0 | 4 | 0 | 1 | 7 |
| 4 | prod | 3 | 6 | 1 | 5 | 0 | 6 | 2 | 11 | 34 |
| 5 | prod | 1 | 0 | 1 | 0 | 0 | 2 | 0 | 1 | 5 |
| 6 | prod | 5 | 1 | 0 | 1 | 0 | 7 | 0 | 6 | 20 |
| 9 | prod | 2 | 1 | 0 | 1 | 2 | 1 | 0 | 2 | 9 |
| 10 | prod | 2 | 0 | 1 | 4 | 2 | 7 | 0 | 1 | 17 |
| 11 | prod | 10 | 7 | 4 | 3 | 2 | 13 | 2 | 10 | 51 |
| 12 | prod | 0 | 2 | 2 | 1 | 0 | 2 | 2 | 4 | 13 |
| 1 | comp | 0 | 2 | 2 | 0 | 0 | 4 | 0 | 3 | 11 |
| 2 | comp | 1 | 0 | 0 | 1 | 0 | 1 | 0 | 5 | 8 |
| 3 | comp | 2 | 4 | 0 | 1 | 0 | 4 | 2 | 4 | 17 |
| 4 | comp | 1 | 5 | 1 | 1 | 0 | 2 | 0 | 9 | 19 |
| 5 | comp | 1 | 3 | 0 | 0 | 0 | 1 | 1 | 1 | 7 |
| 6 | comp | 0 | 0 | 0 | 0 | 0 | 0 | 0 | 2 | 2 |
| 9 | comp | 5 | 4 | 1 | 4 | 5 | 2 | 1 | 3 | 25 |
| 10 | comp | 3 | 0 | 2 | 3 | 5 | 4 | 0 | 0 | 17 |
| 11 | comp | 0 | 1 | 1 | 1 | 0 | 3 | 1 | 9 | 16 |
| 12 | comp | 1 | 4 | 1 | 0 | 0 | 1 | 0 | 4 | 11 |
| total: |  | 40 | 45 | 21 | 33 | 17 | 80 | 15 | 85 | 336 |

